# Modelling shiftwork and sleep fragmentation in mice: behavioural phenotypes and implications for hippocampal neurogenesis

**DOI:** 10.64898/2026.09.26.754534

**Authors:** Robert M Gibson, Taha Yildirim, Kim Simon, Jefferey Yue, Sean Tok, Ralph E Mistlberger, Brianne A Kent

**Affiliations:** Department of Psychology, Simon Fraser University.; Department of Molecular Biology and Biochemistry, Simon Fraser University.; Institute for Neuroscience and Neurotechnology, Simon Fraser University.

**Keywords:** sleep, hippocampal neurogenesis, shiftwork, sleep fragmentation, mice, corticosterone

## Abstract

Sleep disturbances compromise cognition, and overall physical and mental health; however, animal models of these effects remain limited, and typically impose more extreme disturbances than those experienced by humans. Here, we use behavioural interventions in mice to model two common types of sleep disruption: (1) sleep displacement associated with shiftwork and (2) sleep fragmentation associated with aging and disease. C57BL6/J mice were subjected to either a sleep deprivation protocol (*n* = 8, females = 4) or a fragmentation protocol (*n* = 8, females = 4), in which they were completely deprived of sleep or awoken every 2 min respectively, for 12 hours during their typical rest phase (lights-on) across 7 consecutive days. Age-matched control mice (*n* = 14, females = 7) remained undisturbed in their home cages. Activity during the 12-hour active phase was recorded via infrared motion sensors. After protocol completion, mice were injected with bromodeoxyuridine (BrdU, 200mg/kg) 2 hours prior to brain and blood collection; hippocampal neurogenesis and plasma corticosterone were then assessed. Results show that the sleep deprivation and fragmentation protocols had different, dynamic effects on behaviour; sleep deprivation was associated with an accumulating sleep need, operationalized as progressively reduced locomotor activity across the week, whereas fragmentation produced only an acute reduction of locomotor activity. Cell proliferation rates and corticosterone did not differ from controls. These protocols provide a foundation for future research into the mechanistic links between shiftwork and sleep fragmentation on health and cognition.

## Introduction

Sleep is fundamentally important for optimal cognitive functioning, underpinning critical processes such as memory consolidation, attention, and executive function (Hyndych et al., 2025; Rasch & Born, 2013). Acute and chronic sleep disruption produce measurable deficits across a broad spectrum of neuropsychological domains (Bubu et al., 2017; Rasch & Born, 2013; Zamore & Veasey, 2022). Common causes of sleep disruption include sleep displacement from shiftwork, jetlag, or social jetlag, and sleep fragmentation resulting from aging or disease.

Shiftwork is broadly defined as working outside of typical daytime hours at least periodically and accounts for almost a third of workers in the US (Alterman et al., 2013; Wong et al., 2019). Shift workers often experience poor sleep quality, reduced sleep duration, increased insomnia, circadian misalignment, and difficulty maintaining wakefulness during active periods (Boivin et al., 2022). Night work in particular is associated with acute and chronic deficits in both memory and attentional domains (Boivin et al., 2022; Hudson et al., 2020). Animal models of shiftwork have demonstrated circadian disruption, evident in behavioural, metabolic and circadian clock gene rhythms (Barclay et al., 2015), with adverse effects on metabolic and cognition functions (Hairston et al., 2005).

Sleep fragmentation is a sleep disturbance characterized by frequent interruptions, resulting in decreased consolidation of sleep bouts and sleep quality (McCarter et al., 2022). Fragmentation of sleep is common in older adults and is associated with cognitive decline as well as alterations in brain structure (reviewed by Baillet & Schmidt, 2020). Sleep fragmentation is also common in neurological disorders such as Alzheimer’s disease (AD) and may accelerate the development of underlying neuropathology (reviewed by Kent et al., 2021). Rodent models of chronic sleep fragmentation demonstrate reduced dendritic arborization within regions of the hippocampus (Konakanchi et al., 2022) and spatial memory impairments (Sportiche et al., 2010).

One mechanism implicated in the relationship between sleep-wake cycle disturbance and impairments in cognition is hippocampal neurogenesis (Aimone et al., 2014; Kent & Mistlberger, 2017). Hippocampal neurogenesis encapsulates the proliferation, differentiation, maturation, and survival of adult-born neurons within the granule cell layer of the dentate gyrus, supporting brain plasticity, hippocampal-dependent memory, and resilience to cognitive decline (Aimone et al., 2014; Montaron et al., 2020). Adult-born neurons play an important role in pattern separation, a computational mechanism engaged during encoding that generates distinct neural representations from overlapping inputs, thereby minimizing interference between similar memories (Bekinschtein et al., 2014; Oomen et al., 2014).

Extensive and continuous disruption of sleep-wake cycles (up to 4 days) in rodents utilizing total sleep deprivation, sleep restriction, and fragmentation paradigms have demonstrated reductions in both proliferation and survival rates of adult-born neurons (Fernandes et al., 2015; Guzmán-Marín et al., 2003; Guzmán-Marín et al., 2005; Guzmán-Marín et al., 2007; Mueller et al., 2008; Sompol et al., 2011). However, acute sleep manipulations (24 hours or less) result in decreases, increases, or no effect on neurogenesis (Zucconi et al., 2006; Junek et al., 2010; Roman et al., 2005; van der Borght et al., 2006). Conflicting findings may reflect a dose-response relationship, where brief sleep loss has no effect or even acutely increases neurogenesis, whereas long-term disruptions markedly reduce neurogenesis. However, much of this work uses methodologies that reduce total sleep time to extreme levels (e.g., 7% across 96h Guzmán-Marín et al., 2003, 2005), far beyond what is typically experienced in humans even with significant sleep disorders. Thus, to better understand the effects of sleep disruption on cognition in humans, there is a need for translationally relevant animal models.

In this study, we developed models of shift work and sleep fragmentation in young adult mice to evaluate behavioural and neurogenic consequences. Our manipulations were applied for 12 hours daily, allowing for recovery sleep during the active phase to better represent the intermittent and partial sleep loss experienced by shift workers and individuals with fragmented sleep. We hypothesized that both models would disrupt locomotor activity rhythms and increase sleep pressure, with effects compounding over consecutive days of manipulation. We used gentle handling to minimize effects of sleep manipulations on corticosterone levels, aiming to provide a model for isolating the impact of sleep disruption on neurogenesis from stress-related confounds. We hypothesized that both models would reduce the proliferation of adult-born neurons in the hippocampus compared to age-matched controls.

## Methods

### Animals

All experimental protocols were approved by the Simon Fraser University Animal Care and Use Committee (Protocol #1325P-21). C57Bl6/J mice (*N* = 30; Charles River Laboratories, Montreal, Quebec) were assigned to 1 of 4 experimental conditions: a sleep deprivation (SD) group (*n* = 8) modelling night shift work, a sleep fragmentation (SF) group (*n* = 8), and two respective control groups (SD control *n* = 6, SF control *n* = 8). The SD group and respective controls were 8 weeks old at the beginning of manipulations while the SF group and respective controls were 11 weeks old.

### Housing and Handling Procedures

All animals were housed on a 12:12 light-dark (LD) cycle with lights on (zeitgeber time, ZT 0) at 9am PST for all experimental animals and 9pm PST for all control animals. The opposite LD cycles were used for experimental and control groups to ensure that noise during manipulations occurred during the control group’s active phase, minimizing sleep disturbance for controls. All animals were individually housed in light-proof and sound attenuating cabinets equipped with overhead infrared sensors (Sparkfun Electronics, CO, USA) integrated with Clocklab Data Acquisition Software (Actimetrics, IL, USA) providing counts of activity across each minute of the day. Mice were all put on respective light cycles at least three weeks before commencing sleep disruption procedures. All mice were kept on *adlib* food throughout experimentation and weighed at lights on (ZT 0) each day during manipulations to ensure bodyweight was maintained.

### Sleep Deprivation and Fragmentation Procedures

Sleep manipulation occurred during ZT0-ZT12 for 7 consecutive days. Mice were kept awake by gentle paintbrush prodding or handling in response to behavioural signs of sleep (e.g., curled posture, inactivity, closed eyes). In the SD group, all sleep was prevented, whereas in the SF group mice were disturbed at 2-min intervals when showing signs of sleep, the same interval used by automated devices described in Ramesh et al. (2009). If mice in either group were not showing behavioural signs of sleep, they were left undisturbed but closely monitored.

### Bromodeoxyuridine (BrdU) Administration

On the 7^th^ day of sleep manipulations, experimental mice were administered 200mg/kg BrdU (Sigma-Aldrich, St. Louis, MO, USA, Cas# 59-14-3) intraperitoneally (IP, 10mg/mL in saline) at ZT10, 2 hours prior to the completion of manipulations and euthanasia. Control animals were injected at a matched circadian time.

### Tissue Collection

Immediately upon completion of the 7^th^ day of manipulation, mice were anesthetized (5% isoflurane) and euthanized with CO2. Control animals were euthanized using the same methods at a matched circadian time (ZT 12). Blood samples were collected via cardiac puncture using a 23G needle from the right atrium of the heart, then mixed with 15uL of ethylenediaminetetraacetic acid (EDTA, 100 mg/mL) to prevent coagulation. The elapsed time between euthanasia and blood collection was less than 3 min for all animals. Blood samples were centrifuged (1500g, 20 min, 4°C) after which the supernatant was collected and stored at −80°C. Following cardiac puncture, mice were transcardially perfused with ice-cold Phosphate-Buffered Saline (PBS, pH 7.4) and brains were extracted. Whole brains were stored for 16 h in paraformaldehyde (PFA) at 4°C, then transferred to a 30% sucrose solution until saturated. Brains were sectioned at 40µm using a cryostat and then stored in an antifreeze PBS solution (30% Sucrose, 30% Ethylene Glycol, 0.1% Sodium Azide, 1% Polyvinylpyrrolidone, 1X PBS). Forty-eight sections were collected for each mouse spanning 1.92mm of hippocampal tissue and stored at -20°C.

### Behavioural Analysis

Analysis of locomotor activity was conducted using Clocklab Analysis software (version 6.0.53; Actimetrics, IN, USA). To assess the behavioural and circadian effects of sleep manipulations, we compared locomotor activity during the active phase on experimental days to the average of weeklong baseline measures. To control for variability across sensors, all measures were converted to a percent change relative to each individual animal’s weeklong baseline averages. To evaluate the effects of protocols on behavioural state, a score of ‘wake’ was assigned to every 1 min time bin in which one or more activity counts were registered, and ‘rest’ to 1-min bins with zero counts. We calculated percent change in total wake time (sum of all 1-min bins with at least 1 count of activity), total activity (sum of all counts), and average rest bout length (consecutive 1-min bins with no activity) across each day of experimentation. To assess the effects of the manipulations across circadian phases, average difference waveforms were created by calculating the percentage change in total activity each hour compared to baseline averages for that hour.

### Histology

Six hippocampal sections from each mouse were chosen for free-floating fluorescent immunohistochemistry. Every fifth section (200µm apart) was selected from AP -1.155 mm to -2.255 with exceptions made based on matching of morphological aspects and exclusion of damaged tissues. Sections were treated with 2M HCL at 37°C for 30 min for antigen reveal, incubated in primary antibody Rat anti-BrdU (BU1/75, Abcam Cat# ab6326, RRID:AB_305426) 1:500 in 0.05% 1X Phosphate-Buffered Saline with Tween 20 (PBST) and 2% Normal Goat Serum (NGS) at 4°C for 16 h, secondary antibody Goat anti-Rat (AF-488, Invitrogen) 1:500 in 0.05% 1X PBST and 2% NGS at room temperature for 4 h, and 4′,6-diamidino-2-phenylindole (DAPI) 1:10000 (0.1µg/mL) in 0.05% 1X PBST for 15 min.

### Confocal Imaging

Sections were imaged using a Nikon A1R Laser Scanning Confocal system at 20x magnification (DAPI λ = 404.4, FITC λ = 488). Imaging was performed at 5 different z-levels 7µm apart covering a range of 28µm total. Images from each z-level were then combined into one max-intensity image which was exported for cell counting analysis. The left and right dentate gyri were imaged and analyzed independently, resulting in 12 images per mouse.

### Cell Quantification and Analysis

Images were analyzed using ImageJ (version 1.54f). All images were cropped to isolate the dentate gyrus, then brightness and contrast were adjusted using the auto function (equivalent to: run "Enhance Contrast", "saturated=0.35"). Images were split into two monochromatic channels (DAPI and FITC) and the granule cell layer (GCL) was identified as a region of interest (ROI) which was then extended by 15 pixels (19.14µm) to capture the subgranular zone (SGZ), similar to the approach described in Mueller et al. (2008). To distinguish proliferating cells from artifact we excluded FITC signal within this ROI based on size (minimum = 20 pixels/12.76µm, max = 150 pixels/191.4µm), intensity (minimum = 59), and circularity (minimum = 0.4). Then each image was evaluated independently by three trained researchers, blind to experimental conditions, to confirm the accuracy of both the area of the GCL and validity of each individual FITC signal. Furthermore, to assess density of proliferation, the area estimates of GCL and SGZ were first converted from pixels to um^2^, then multiplied by 40um (section size), and converted from um^3^ to mm^3^ to get estimates of the volumetric space analyzed. Inter-rater reliability was high (>98% agreeance) for both raw cell counts as well as area estimates of the GCL and SGZ, thus the averages across scorers were used.

### Corticosterone ELISA

The level of corticosterone was measured using the DetectX® Corticosterone Enzyme Immunoassay Kit (Arbor Assays, Ann Arbor, MI, USA) following the manufacturer’s instructions. A standard curve was generated using a four-parameter logistic curve (4PLC) fitting routine. The antiserum cross-reacts 100% for corticosterone, 18.90% for 1-dehydrocorticosterone, 12.30% for deoxycorticosterone, 3.30% for 1α-hydroxycorticosterone, and 2.44% for 11-dehydrocorticosterone. The intra-assay coefficient of variation was calculated as 15.53%.

### Data Analysis

All analyses were conducted using GraphPad Prism (v.11, GraphPad Software, Boston, Massachusetts USA). The 1.5x interquartile range (IQR) rule was used to exclude extreme values except for analyses including sex as a factor due to sample size limitations.

## Results

### Sleep deprivation and fragmentation produce differentiable, dynamic behavioural phenotypes

Behavioural effects of SD included significant main effects across days for total wake time (mixed-effects ANOVA, *F*(2.699, 17.1) = 39.82, p < 0.0001; Fig 1A), total activity (*F*(2.306, 14.99) = 16.62, p = 0.0001; Fig 1D), and average rest bout length (*F*(2.639, 18.03) = 29.2, p <0 .0001; Fig 1G). Behavioural effects of SF included significant main effects of day on total wake time (*F*(3.073, 21) = 9.745, p = 0.0003; Fig 1B), total activity (*F*(2.722, 17.69) = 5.667, p = 0.0078; Fig 1E), and average rest bout length (*F*(1.967, 13.44) = 11.1, p = 0.0015; Fig 1H).

**Figure 1:**
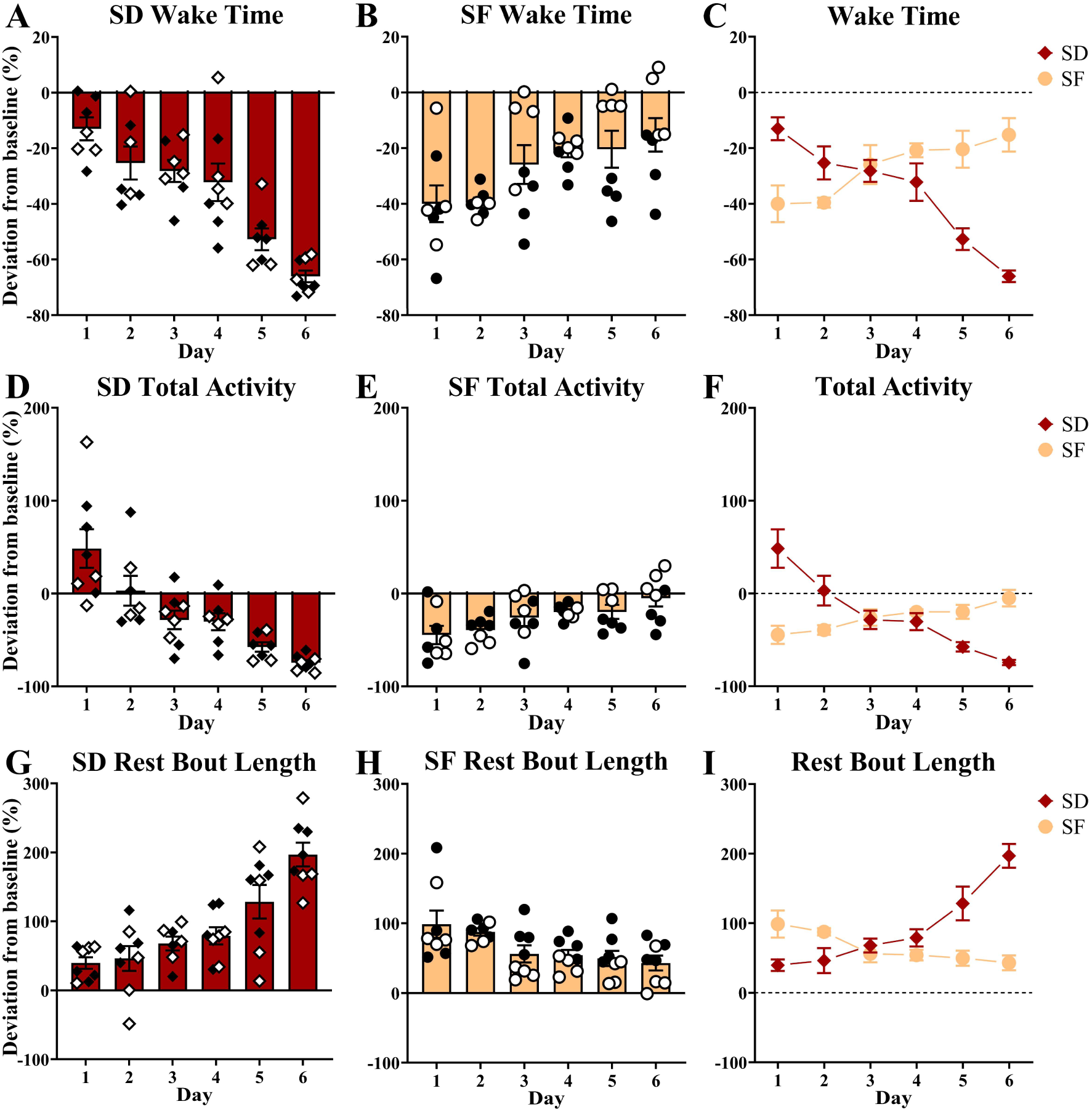
Behavioural effects of SD and SF protocols. Deviation from baseline in active phase wake time across days during **(A)** SD, **(B)** SF, and **(C)** a comparison between the two protocols. Deviation from baseline in active phase total activity during **(D)** SD, **(E) SF**, and **(F)** a comparison between the two protocols. Deviation from baseline in the average active phase rest length duration during **(G)** SD, **(H)** SF, and **(I)** a comparison between the two protocols. Data expressed as mean +/- SEM. SD, sleep deprivation shown in red; SF, sleep fragmentation shown in orange. Open data points represent females. Hollow data points represent females

To investigate the differential behavioural effects of the protocols, we compared the percent change in time awake from baseline across days and found a significant main effect of day (mixed-effect ANOVA, *F*(3.846, 50) = 4.062, p = 0.0069; Fig 1C), no main effect of protocol (*F*(1,14) = 2.949, p = 0.108), and a significant interaction between day and protocol (*F*(3.846, 50) = 23.82, p < 0.0001). A Šidák multiple comparisons test indicated that on day 1 SF reduced daily wake time compared to SD (*t*(11.44) = 3.471, p = 0.0293); however, on day 5 (*t*(11.13) = 4.185, p = 0.0089) and day 6 (*t*(8.676) = 7.983, p = 0.0002) SD reduced wake time significantly more than SF. Furthermore, the change in total activity revealed a significant main effect of day (mixed-effect ANOVA, *F*(2.734, 34.99) = 5.562, p = 0.004; Fig 1F), no main effect of protocol (*F*(1, 14) = 0.1896, p = 0.6688), and a significant interaction between day and protocol (*F*(2.734, 34.99) = 20.14, p < 0.0001). A Šidák post-hoc analysis revealed that SF produced a larger decrease in activity counts on day 1 (*t*(9.994) = 4.052, p = 0.0138) while SD resulted in a greater reduction on day 5 (*t*(10.55) = 4.131, p = 0.0109) and day 6 (*t*(8.431) = 7.422, p = 0.0003). Finally, the change in average rest bout length showed a significant main effect of day (mixed-effects ANOVA, *F*(3.364, 45.76) = 7.973, p = 0.0001: Fig I), no significant main effect of protocol (*F*(1, 14) = 4.191, p = 0.0599), and a significant interaction between day and protocol (*F*(3.364, 45.76) = 24.6, p < 0.0001). Šidák post hoc tests reveal that on day 6 SD produced a significantly greater increase (*t*(11.71) = 7.629, p < 0.0001) in average rest bout length compared to SF.

### Sleep disruption protocols did not affect cell proliferation

Effects of sleep manipulations on cell proliferation were analyzed independently for SD and SF groups, to avoid potential confound effects of age. Of the 336 images, 25 were removed from analysis including all 12 from a single SD control female due to an incomplete BrdU injection, as well as 13 others due to tissue damage or artifact. First, we quantified cell proliferation as the number of BrdU+ cells per ROI, averaged across sections for each mouse, and there were no differences between SD and the controls (*t*(8) = 0.8043, p = 0.4445; Fig 2A), or SF and their respective controls (*t*(14) = 0.8203, p = 0.4258; Fig 2B). To control for the effects of individual differences in DG morphology, we next assessed the density of BrdU+ cells by dividing the raw count of BrdU+ cells in each section by the volume of the ROI (GCL + SGZ) analyzed. Neither SD (*t*(7) = 1.218, p = 0.2626; Fig 2C), nor SF (*t*(12) = 0.3793, p = 0.7111: Fig 2D), affected the density of BrdU+ cells compared to controls. Notably, our data highlight a potential sex-dependent effect of SF as the density of BrdU+ cells increased in males compared to same-sex controls but in females decreased along with BrdU+ cells/section (Supplementary Figure S2). However, this finding will need to be substantiated in a larger sample.

**Figure 2:**
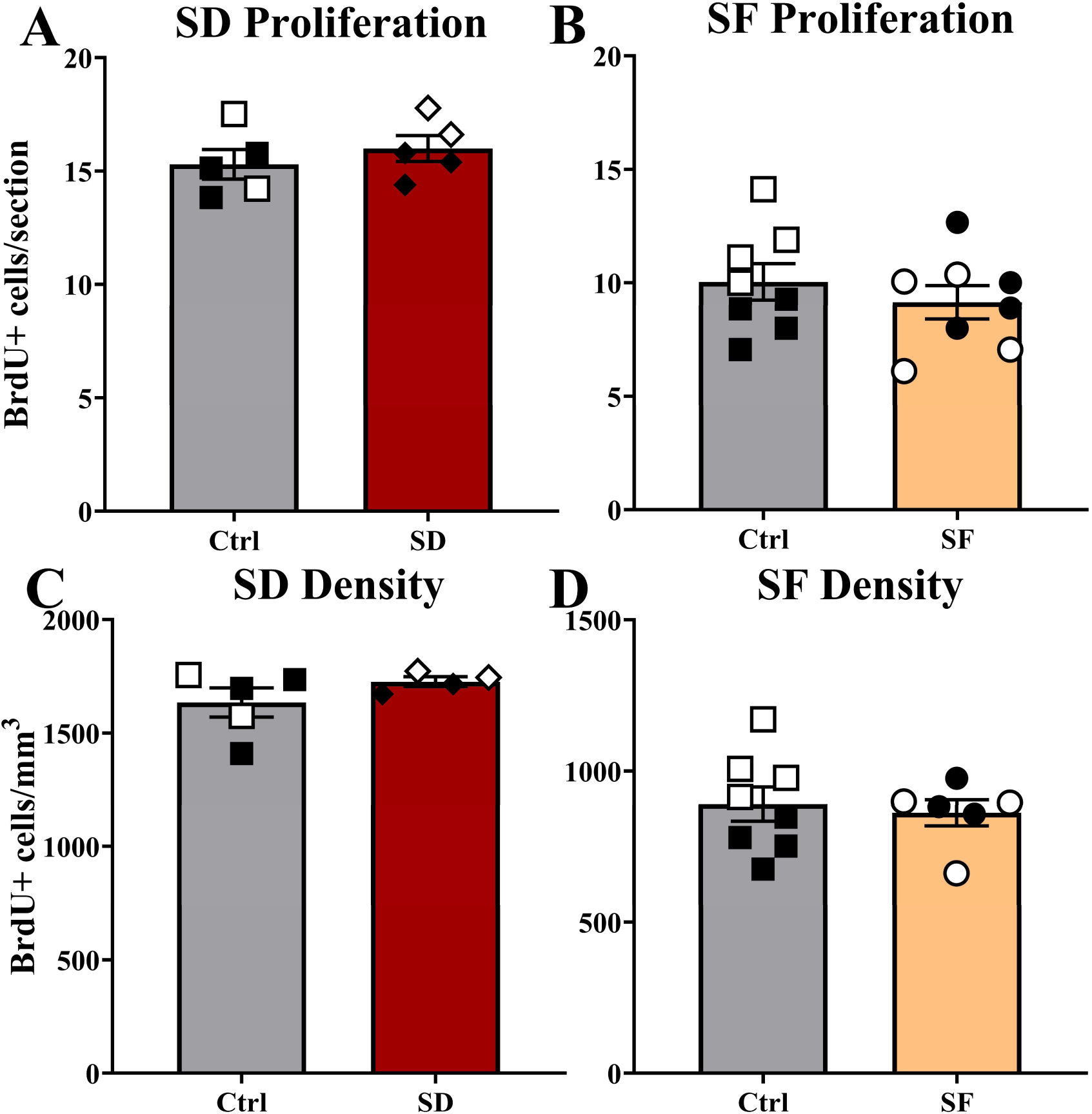
Effects of protocols on hippocampal neurogenesis. **(A)** SD and **(B)** SF effects on cell proliferation in the dentate gyrus (DG). **(C)** SD and **(D)** SF effects on proliferation density in the DG. Data expressed as mean +/- SEM. Hollow data points represent females. BrdU, bromodeoxyuridine; Ctrl, control group; SD, sleep deprivation; SF, sleep fragmentation.

### Sleep disruption protocols did not affect plasma corticosterone concentration

To assess whether our experimental manipulations induced stress, a known modifier of neurogenesis, we analyzed corticosterone levels. Neither SD (*t*(10) = 0.4583, p = 0.6565: Fig 3A) nor SF (*t*(13) = 0.034, p = 0.9734; Fig 3B) altered corticosterone concentrations compared to respective control groups.

**Figure 3:**
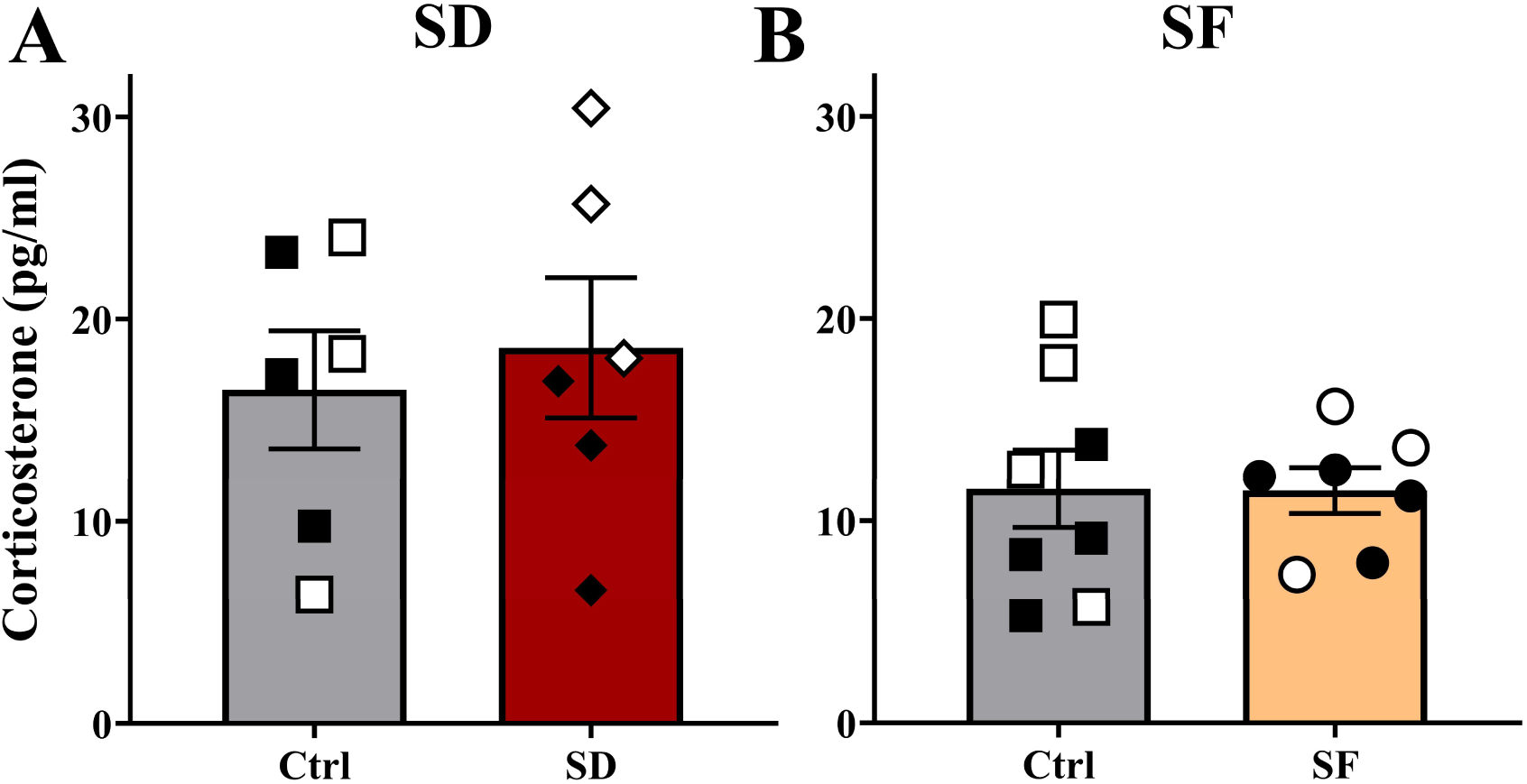
Effects of sleep manipulations on corticosterone. **(A)** SD and **(B)** SF effects on corticosterone levels (pg/ml) compared to their respective controls. Data expressed as mean +/- SEM. Hollow data points represent females. Ctrl, control group; SD, sleep deprivation; SF, sleep fragmentation.

## Discussion

Sleep-wake cycle disruptions, such as with shiftwork and the sleep fragmentation common with aging or disease, negatively affect cognition. We modelled shiftwork, using a sleep deprivation protocol, and sleep fragmentation to evaluate the effects on activity, hippocampal neurogenesis, and corticosterone levels in mice. Unexpectedly, the two protocols produced distinct, dynamic behavioural effects rather than differing simply in severity: fragmentation drove a more pronounced acute reduction in locomotor activity, suggestive of an immediate sleep rebound, while the shiftwork model produced an accumulating effect across the week.

Specifically, the behavioural phenotype caused by the shiftwork model was suggestive of progressively increasing sleep debt: during the active phase, wake duration and total activity decreased while average rest bout length increased. This behavioural pattern recapitulates daytime sleepiness, aligning with clinical presentations of sleep-wake cycle disruption in humans (Boivin et al., 2022). The decrease in total activity aligns with previous work by Barclay et al. (2014) which demonstrated that depriving young adult C57Bl6/J mice of sleep during the first 6 hours of their light phase for 5 days a week, decreased locomotor activity during the active phase and disrupted both molecular and metabolic circadian rhythms.

In contrast, the sleep fragmentation model produced an acute decrease in wake duration and total activity after a single day of manipulation, suggesting a rapid increase in sleepiness. Activity levels then gradually returned to baseline, suggesting that animals accommodated to the procedure across successive days. To our knowledge, this is the first study to specifically assess the dynamic changes in active phase locomotor activity during weeklong sleep fragmentation in mice. Future work should assess whether this dynamic response is also seen in humans experiencing sleep fragmentation in both acute and chronic contexts.

The dynamic behavioural effects produced by the two models were distinguishable from one another. At manipulation onset, only the fragmentation paradigm acutely decreased locomotor activity; the shiftwork model did not. After multiple consecutive days, the fragmentation paradigm’s effects on arousal normalized, whereas the shiftwork protocol produced a significant decrease in arousal, suggestive of increasing sleep need. This divergence highlights the value of using the same stimulus, applied differently, to model distinct human sleep disruptions. The increasing deviation from baseline across successive days seen in our model of shiftwork also suggests that effects may continue to increase with extended duration. Future research exploring whether this same pattern holds true across longer disruption periods is needed to better understand the threshold at which neurogenesis is affected.

Despite the significant behavioural effects of the protocols, our analyses revealed no effect on cell proliferation in the hippocampus. This aligns with previous research implementing acute sleep deprivation (12h beginning at ZT0) for a single day showing negligible effects on neurogenesis or even increases (Zucconi et al., 2006; Junek et al., 2010; van der Borght et al., 2006). Our results suggest that when given adequate recovery sleep, even at an inappropriate circadian time, weeklong sleep disruption does not reduce levels of cell proliferation. Previous work by Guzmán-Marín et al. (2007) and Sportiche et al. (2010) has demonstrated that prolonged sleep fragmentation reduced hippocampal neurogenesis, however, our work suggests that shorter periods across consecutive days may not reach the threshold of disruption at which neurogenesis is affected. Given the role of neurogenesis in memory (Bekinschtein et al., 2014; Clelland et al., 2009; Hairston et al., 2005; McTighe et al., 2009) and established relationship between sleep disruptions and physiological brain changes in aging and neurodegenerative disorders such as AD (Baillet & Schmidt, 2020; Bubu et al., 2017; Kent & Mistlberger, 2017; Lim et al 2013; Mattis & Sehgal, 2016; Peter-Derex et al., 2015; Shi et al., 2018; Wang & Holtzman, 2020; Wang et al., 2024) further investigation into this potential threshold using translational pre-clinical models is merited.

Our analyses of proliferation focused on the cumulative effects of the weeklong sleep disruption paradigms; this approach may have missed acute effects on neurogenesis that were attenuated by recovery sleep. We also assessed proliferation only, and did not directly evaluate maturation, differentiation, nor survival of adult-born neurons, all of which may be affected by sleep disruption in rodents (Zucconi et al., 2006; Guzmán-Marín et al., 2005, 2007; Hairston et al., 2005; Mueller et al., 2008; Sportiche et al., 2010). Additionally, our model lacks control for the locomotion induced by sleep disruption. Exercise in running wheels is well established to increase hippocampal neurogenesis (Ben-Zeev et al., 2022; Kent et al., 2015; van Praag, 2008), but the amount of locomotor activity associated with the sleep disruption procedures used in this study is trivial by comparison. Finally, despite the translational appeal of our weeklong procedures compared to existing models, our manipulations may be too brief to capture the health and neurobiological consequences of long-term shiftwork and sleep fragmentation.

A strength of our models is that they are less stressful than other existing methods. Plasma corticosterone levels suggest that neither manipulation increased stress compared to respective controls. This is important, as sleep disruption protocols are inherently stressful, and stress hormones reduce hippocampal neurogenesis (Gould et al., 1997; Toda et al., 2019). We interrupted sleep using gentle handling upon behavioural signs of sleep onset, rather than automated methods (ie. treadmill, rotating bar, etc.), increasing the translational potential of our findings. Thus, our methodology can be used in future work to assess hippocampal neurogenesis with a limited impact on stress hormones.

## Conclusion

Weeklong models of shiftwork and sleep fragmentation produce distinguishable, dynamic behavioural phenotypes in rodents that recapitulate rest-activity disruptions observed in humans. The temporal dissociation between these phenotypes offers a novel tool for identifying the thresholds at which specific patterns of sleep disruption produce neurobiological and cognitive consequences — insight that could inform mechanisms of disease and targets for therapeutic intervention.

## Acknowledgements

Afnan Sahibzada, Manthan Vekariya, Mateusz Michalik, Mayuko Arai, Emad Shams, Grace Budvardson, Victoria Carriquiriborde, Kashish Mehta, Arman Virk, Dylan Yee & Luis Munoz.

## Data availability statement

The data will be openly available upon publication using the SFU Institute for Neuroscience and Neurotechnology’s data sharing platform.

