## Supplementary material for "Modelling shiftwork and sleep fragmentation in mice: behavioural phenotypes and implications for hippocampal neurogenesis"

#### Difference Waveforms

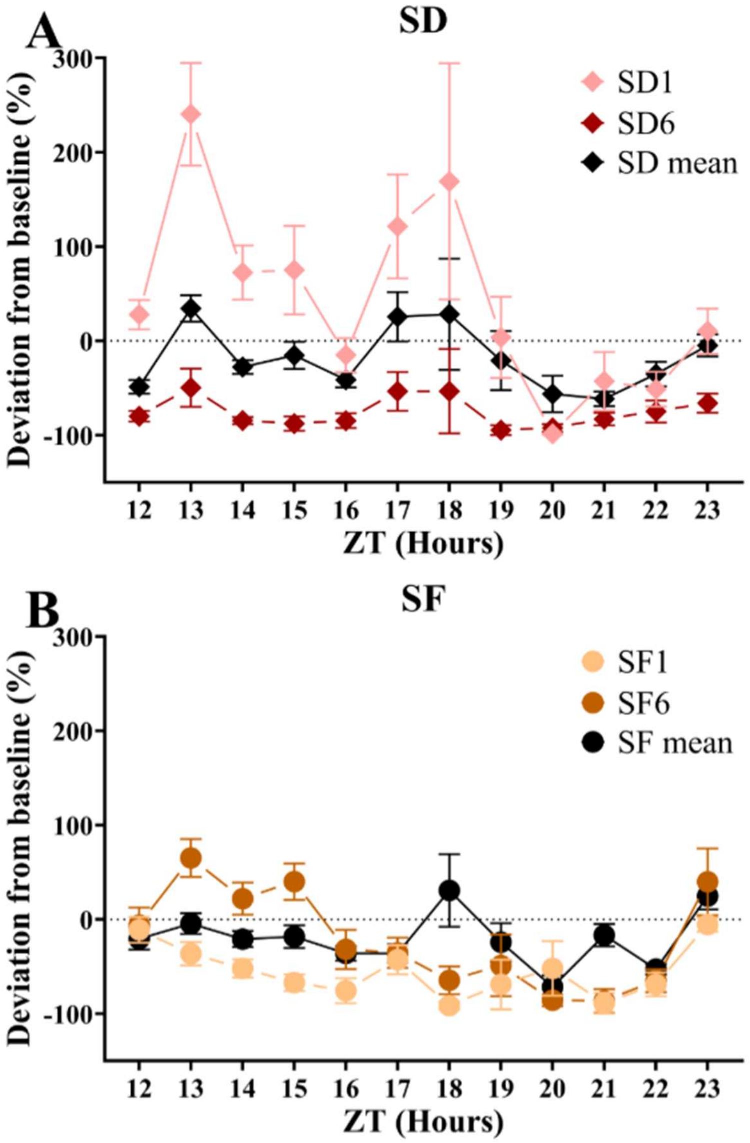


**Figure S1:** Effects of sleep manipulations on daily rhythms of activity. Difference between total activity at each hour following first day of manipulation (SD1 and SF1), sixth day of manipulation (SD6 and SF6), and mean across the 6 days compared to baseline during **(A)** SD and **(B)** SF. Data expressed as mean +/- SEM. SD, sleep deprivation; SF, sleep fragmentation; ZT, zeitgeber time.

Analysis of the average difference waveform for SD revealed a significant main effect of ZT (mixed-effects analysis, $F(1.931, 13.52) = 4.826$, $p = 0.0271$), day ($F(1.129, 7.901) = 26.77$, $p = 0.0007$) and a significant interaction ($F(2.946, 19.01) = 3.818$, $p = 0.0274$). A Šídák's multiple comparisons test showed a significantly more positive deviation from baseline on day 1 compared to both day 6 ($t(7) = 7.252$, $p = 0.0061$) and the average across SD ($t(7) = 7.376$, $p = 0.0055$) at ZT12 as well as a significantly more negative deviation from baseline on day 6 compared to the average across SD ($t(5) = 6.888$, $p = 0.0349$) at ZT14. Analysis of the average difference waveform for SF revealed a main effect of ZT (mixed-effects analysis, $F(3.825, 26.78) = 6.372$, $p = 0.0011$), day ($F(1.935, 13.55) = 14.79$, $p = 0.0004$), and a significant interaction ($F(4.159, 26.65) = 4.026$, $p = 0.0104$). A Šídák's multiple comparisons test showed a significantly more negative deviation from baseline on day 1 compared to the average across SF ($t(6) = 7.016$, $p = 0.015$) at ZT21.

**Cell Proliferation by Manipulation & Sex**

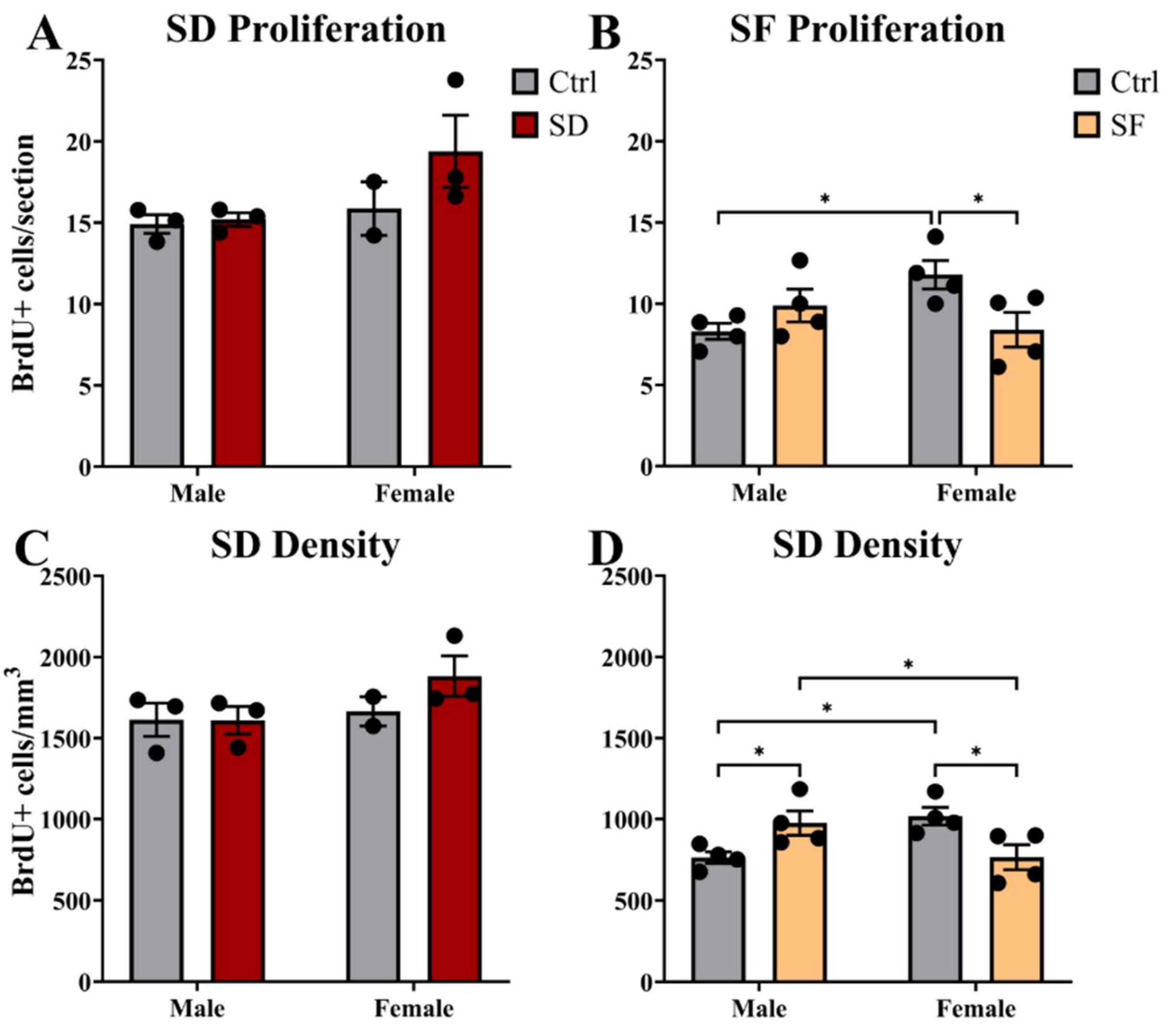


**Figure S2:** Sex dependent effects of sleep manipulations on hippocampal neurogenesis. **(A)** SD and **(B)** SF effects on cell proliferation in the dentate gyrus (DG). **(C)** SD and (**D)** SF effects on proliferation density in the DG . Data expressed as mean +/- SEM. BrdU, bromodeoxyuridine; Ctrl, control group; Exp, experimental group; SD, sleep deprivation; SF, sleep fragmentation; * indicates $p < 0.05$

Given there are documented differences in neurogenesis rates between sexes (Deng et al., 2024; Yagi & Galea, 2019) we assessed the contribution of sex in our models. There was no effect of SD on proliferation or density ($p > 0.05$); however, SF had sex-dependent effects such that there was no significant main effect of protocol (two-way ANOVA, $F(1, 12) = 1.014$, p =

0.3339) or sex ($F(1, 12) = 1.255$, p = 0.2844), but there was a significant interaction ($F(1, 12)$ = 7.835, p = 0.0161). A post-hoc comparison revealed that females exposed to SF had significantly fewer $BrdU^+$ cells/section compared to same-sex controls ($t(12) = 2.691$, p = 0.0196), and that female controls had more compared to male controls ($t(12) = 2.772$, p = 0.0169). A two-way ANOVA assessing the effects of SF on the density of $BrdU^+$ cells revealed no main effect of protocol ($F(1, 12) = 0.1029$, p = 0.7538) or sex ($F(1, 12) = 0.1257$, p = 0.7291) on density, but there was a significant interaction ($F(1, 12) = 13.67$, p = 0.003). A Fisher's LSD test revealed that SF males had a higher density of proliferation compared to male controls ($t(12) = 2.388$, p = 0.0343) and SF females ($t(12) = 2.364$, p = 0.0358), that SF females had lower density compared to female controls ($t(12) = 2.841$, p = 0.0149), and that female controls had higher density than male controls ($t(12) = 2.865$, p = 0.0142).

### Corticosterone by Manipulation & Sex

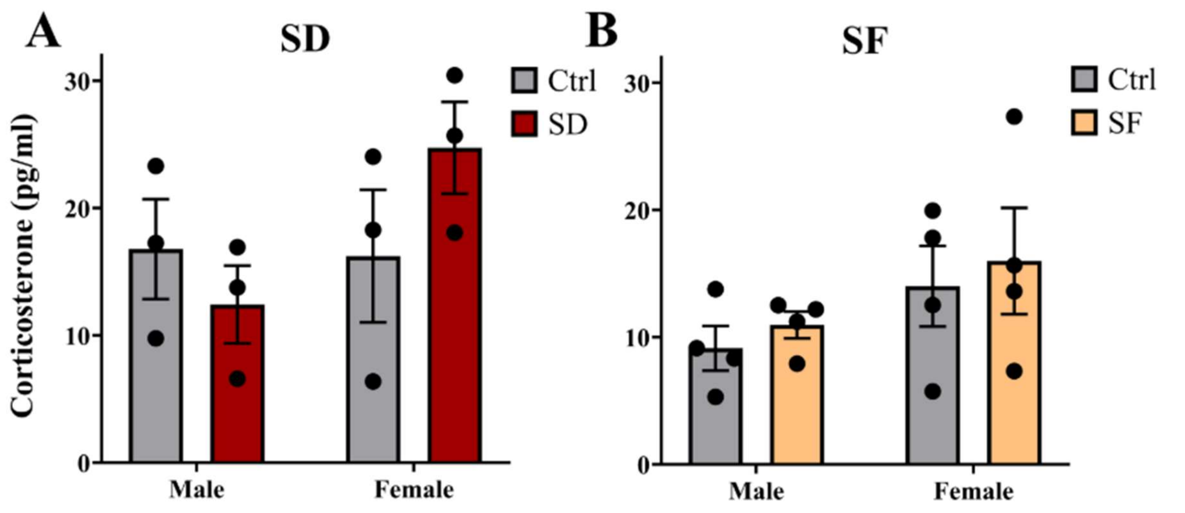


**Figure S3:** Sex dependent effects of sleep manipulations on corticosterone. Corticosterone levels (pg/ml) of mice exposed to SD (**A**) and SF (**B**) compared to their respective controls. SD and SF had no effects on corticosterone

($p > 0.05$). Data expressed as mean +/- SEM. Ctrl, control group; Exp, experimental group; SD, sleep deprivation; SF, sleep fragmentation.

There was no effect of SD nor SF on corticosterone ($p > 0.05$) when sex was included in our analyses.
